# A climosequence of chronosequences in southwestern Australia

**DOI:** 10.1101/113308

**Authors:** Benjamin L. Turner, Patrick E. Hayes, Etienne Laliberté

## Abstract

To examine how climate affects soil development and nutrient availability over long timescales, we studied a series of four long-term chronosequences along a climate gradient in southwestern Australia. Annual rainfall ranged from 533 mm to 1185 mm (water balance from –900 mm to +52 mm) and each chronosequence included Holocene (≤6.5 ka), Middle Pleistocene (120–500 ka), and Early Pleistocene (∼2000 ka) dunes. Vegetation changed markedly along the climosequence, from shrubland at the driest site to *Eucalyptus* forest at the wettest. The carbonate and P content of the parent sand declined along the climosequence, presumably linked to variation in offshore productivity. However, soil development and associated nutrient status followed remarkably consistent patterns along the four chronosequences. Pedogenesis involved decalcification and secondary carbonate precipitation in Holocene soils and leaching of iron oxides from Middle Pleistocene soils, ultimately yielding bleached quartz sands on the oldest soils. Along all chronosequences soil pH and total P declined, while C:P and N:P ratios increased, consistent with the predicted shift from N to P limitation of vegetation during ecosystem development. The expected unimodal pattern of leaf area index was most pronounced along wetter chronosequences, suggesting an influence of climate on the expression of retrogression. The four chronosequences do not appear to span a pedogenic climate threshold, because exchangeable phosphate and base cations declined consistently during long-term pedogenesis. However, the proportion of the total P in organic form was greater along wetter chronosequences. We conclude that soils and nutrient availability on the coastal sand plains of southwestern Australia change consistently during long-term pedogenesis, despite marked variation in modern vegetation and climate. The four chronosequences provide a rare soil-age × climate framework within which to study long-term ecosystem development.

## Introduction

Pedogenesis over thousands to millions of years involves marked changes in nutrient availability (Vitousek, 2004). In particular, nitrogen (N) increases in the early stages of ecosystem development through biological N fixation, whereas phosphorus (P) availability declines continuously as P is lost by leaching at a greater rate than it is replenished by weathering or atmospheric inputs (Walker & Syers, 1976). This drives a long-term shift from N to P limitation of plant biomass and productivity (Vitousek & Farrington, 1997;Wardle *et al.*, 2004; Laliberté *et al.*, 2012), which is reflected in marked changes in the diversity and function of plant and microbial communities (Williamson *et al.*, 2005;Jangid *et al.*, 2013; Laliberté *et al.*, 2013;Zemunik *et al.*, 2015).

Rates of pedogenesis and nutrient depletion influenced markedly by climate (Chadwick & Chorover, 2001; Selmants & Hart, 2010;Feng *et al.*, 2016). Assuming other soil-forming factors are held constant (i.e. topography, parent material, and vegetation), wetter sites are expected to lose rock-derived nutrients at a greater rate than comparable drier sites due to accelerated weathering and greater leaching losses. However, vegetation typically co-varies with climate and might mitigate nutrient loss. For example, wetter sites tend to support greater plant biomass than drier sites, retaining nutrients in the system and reducing the rate of nutrient loss by leaching, at least when precipitation is in balance with potential evapotranspiration (Porder & Chadwick, 2009). In contrast, arid ecosystems lose nutrients more slowly through reduced leaching but support little plant biomass, and therefore have a limited potential to retain nutrients over long timescales. In other words, plants might be able to influence the rate of nutrient loss during pedogenesis depending on the extent to which they retain nutrients in the plant-soil system, which itself depends on climate (Porder & Chadwick, 2009).

Despite the significance of this soil development model to our understanding of terrestrial nutrient cycling, it has been tested only once (Porder & Chadwick, 2009). This is because doing so requires multiple long-term soil chronosequences that are similar to terms of parent material, topography, and time, yet that differ in terms of potential evapotranspiration, precipitation, and water balance. Model systems that meet these strict requirements are exceedingly rare – so far only the well studied Hawaiian Island sequence has been suitable. Porder and Chadwick (2009)studied three basalt lava flows on Hawaii varying from 10,000 to 350,000 years old, with annual precipitation ranging from 500 to 2500 mm. At relatively dry sites (< 750 mm annual rainfall) where evapotranspiration exceeded precipitation, plants slowed nutrient loss, but the effect was small since there was little plant biomass. At intermediate rainfall sites (750–1400 mm) where potential evapotranspiration exceeded precipitation (i.e., negative water balance), plants uplift of nutrients enriched the soil surface with P for at least 350,000 years, effectively compensating for leaching losses. In contrast, at the wettest sites (>1500 mm annual rainfall) where precipitation exceeded evapotranspiration (i.e., positive water balance), leaching losses overwhelmed the capacity of plants to retain nutrients after 350,000 years of pedogenesis, despite greater biomass. Indeed, on much older soils (Oxisols approximately 4 million years old) this ‘high rainfall’ process domain (see below) occurred above about 900 mm of annual rainfall (compared to 1500 mm on younger soils), presumably reflecting long-term depletion of P, reduced productivity, and therefore a reduced capacity of the ecosystem to retain nutrients against leaching losses (Vitousek & Chadwick, 2013a). The influence of plants on nutrient retention therefore appears to be strongest where potential evapotranspiration is roughly in balance with precipitation, although this can be overridden on strongly weathered soils when extreme P limitation reduces the capacity of plants to retain nutrients in the ecosystem.

The rainfall zones described by Porder and Chadwick (2009)for P dynamics reflect the balance between precipitation and evapotranspiration, and correspond to ‘soil process domains’ (Vitousek & Chadwick, 2013a), defined as climate zones within which soils appear to change relatively little across broad variation in rainfall. The domains are separated by pedogenic thresholds, where soil properties vary abruptly across a relatively short variation in rainfall (Chadwick & Chorover, 2001). For basaltic soils on Hawaii subject to about 150,000 years of pedogenesis, the process domains correspond to soils where evapotranspiration exceeds precipitation and carbonate accumulation dominates pedogenesis (low rainfall < 700 mm), a domain where evapotranspiration is slightly greater than precipitation (700 to 1500 mm) and plant uplift and retention of nutrients can compensate for leaching losses, a wetter domain (1500 to 2500 mm) where precipitation exceeds evapotranspiration and pedogenesis is dominated by the accumulation of metal oxides (Chadwick *et al.*, 2003), and a very wet domain (> 2500 mm) where saturation causes iron to be solubilised and lost via reduction and leaching (Chadwick & Chorover, 2001; Vitousek & Chadwick, 2013a). Above the threshold where precipitation exceeds evapotranspiration the soils exhibit a decline in pH, a decline in base cations and base saturation, and an increase in exchangeable Al (Chadwick *et al.*, 2003). These changes are linked to the depletion of primary minerals and leaching at a greater rate than replenished by weathering.

Here we report a study of a series of four long-term (Early Pleistocene) chronosequences along a rainfall gradient in southwestern Australia. Our recent study of soil development along the Jurien Bay chronosequence in Western Australia, corresponding to the driest chronosequence in the present study, shows a characteristic pattern of soil development, with decalcification of young carbonate-rich dunes, formation of a petrocalcic horizon, and leaching of iron oxide coatings on sand grains leaving bleached quartz sands many meters deep (Turner & Laliberté, 2015). These changes correspond with marked depletion of soil P and other nutrients, leading to some of the most infertile soils in the world on the oldest dunes. Here we extend our Jurien Bay work to similar coastal dune deposits extending hundreds of kilometers south along a strong climate gradient, allowing us to identify three additional long-term soil chronosequences with similar aged dunes but contrasting water balance. Our aim was to use this climosequence of chronosequences along the coast of southwestern Australia to examine how climate influences soil development and nutrient availability during long-term pedogenesis.

## Materials and Methods

### Regional overview and description of the chronosequences

A series of dune deposits occur along parallel to the coast of southwestern Australia, running for approximately 400 km from Geraldton in the north to Dunsborough in the south (Fig. 1). Known as the Swan Coastal Plain, the dunes were formed by periodic interglacial sea-level high stands since the Early Pleistocene or Late Pliocene *(i.e.* 2.59 million years ago) (Kendrick *et al.*, 1991). An additional area of dunes along the southern coastline near Pemberton and Northcliffe, the Scott Coastal Plain, is assumed to correspond to the main dune deposits on the Swan Coastal Plain (Playford *et al.*, 1976). The dunes and their associated soils are grouped into three main units according to the underlying parent sand deposits (McArthur & Bettenay, 1974;Playford *et al.*, 1976). The Quindalup dunes of Holocene age (up to 6500 years old) and correspond with the Safety Bay Sand, the Spearwood dunes are of Middle Pleistocene age (120,000 to 500,000 years old) and correspond with the Tamala Limestone, and the Bassendean dunes are Late Pleistocene in age (approximately 2 million years old) and correspond with the Bassendean Sand.

**Figure 1.**
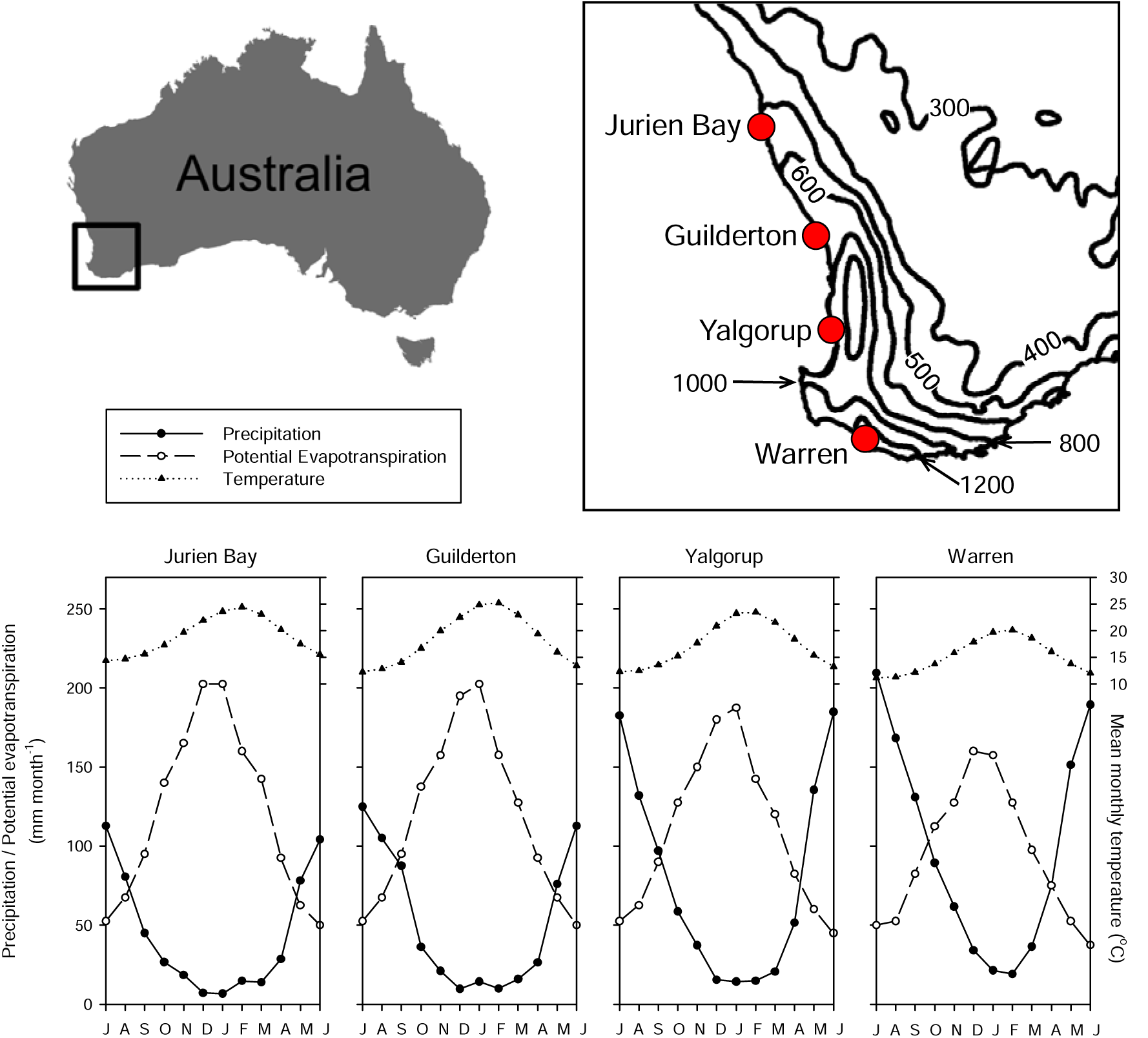
Map of southwestern Australia, showing isolines for annual rainfall (mm year^−1^) and locations of the four dune chronosequences. Rainfall data are averages from the 1961–1990 period (www.bom.gov.au). The lower panel shows monthly climate data, including precipitation, potential evapotranspiration, and mean temperature for each chronosequence.

We previously identified and studied the Jurien Bay chronosequence, located 200 km north of Perth and receiving approximately 530 mm annual rainfall (Fig. 1, 2A) (Laliberté *et al.*, 2012;Hayes *et al.*, 2014;Laliberté *et al.*, 2014; Turner & Laliberté, 2015;Zemunik *et al.*, 2015; 2016). We now identify three additional chronosequences spanning a strong climate gradient along the coastline. These chronosequences are located near Guilderton, approximately 75 km north of Perth (Fig. 2B), Yalgorup National Park, approximately 100 km south of Perth (Fig. 2C), and at Warren Beach, south of Pemberton and approximately 300 km south of Perth (Fig. 2D). Along each of the three new chronosequences we delineated six to seven chronosequence stages: Holocene (stages 1–3), three Middle Pleistocene (stages 4, 5, or 5a and 5b), and one Early Pleistocene (stage 6). In the three drier chronosequences of the Swan Coastal Plain (Jurien Bay, Guilderton, Yalgorup) these stages correspond to the Quindalup (Holocene), Spearwood (Middle Pleistocene) and Bassendean (Early Pleistocene) dunes (Fig. 2A-C) described by McArthur (2004). On the other hand, the first six stages of the Warren chronosequence are classified as Meerup sand or podzols (Fig. 2D), while the oldest stage is mapped as the Cleave series (Purdie *et al.*, 2004).

**Figure 2.**
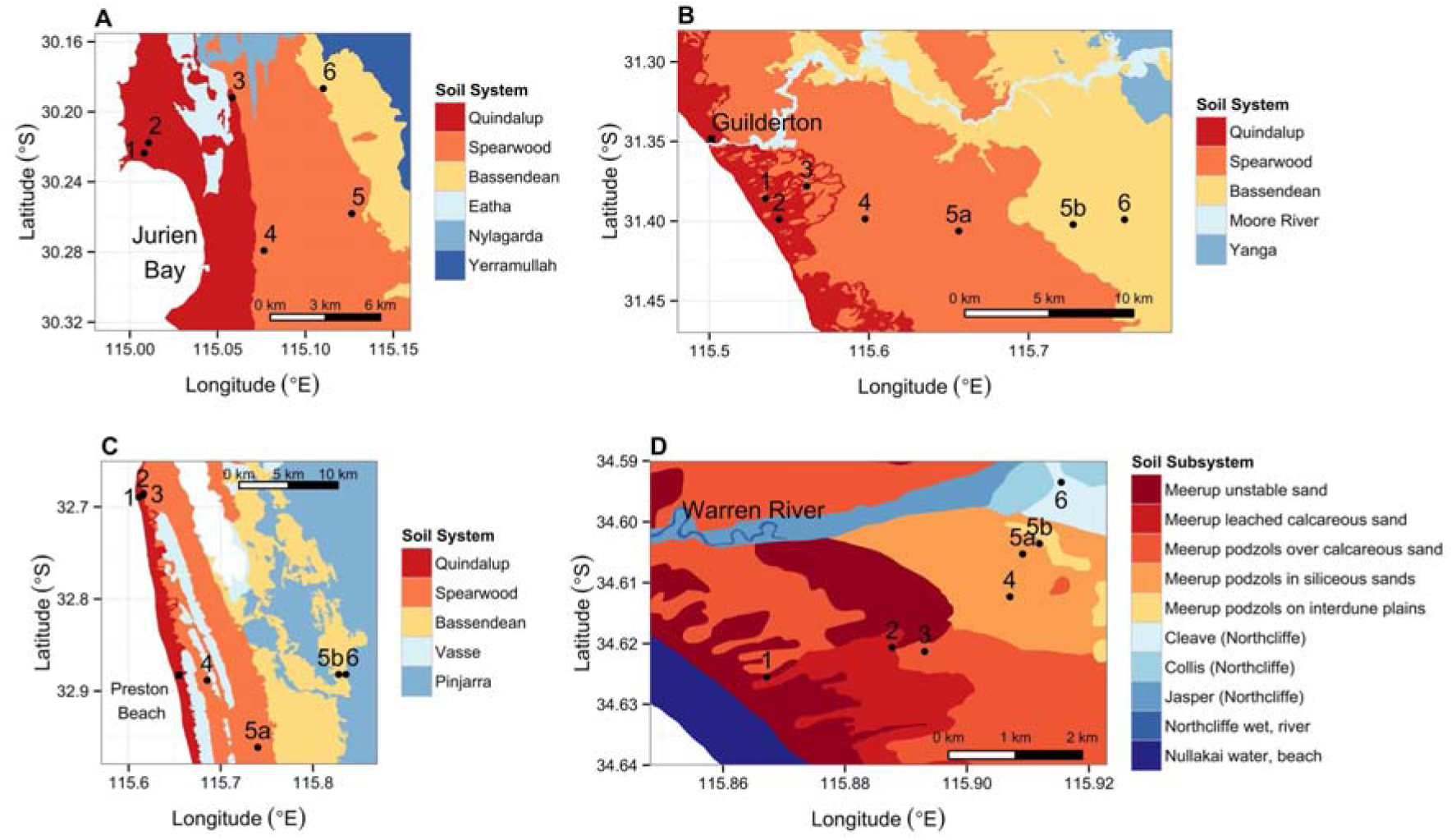
Detailed soil classification maps showing the location of the soil sampling sites along the four dune chronosequences: (A) Jurien Bay, (B) Guilderton, (C) Yalgorup and (D) Warren. Mapping of soil systems and subsystems is based on the Western Australian Department of Agriculture soil classification.

Although precise ages of dune formation are not known for any of the chronosequence stages, various lines of evidence support the chronology ranging from the Holocene to the Early Pleistocene (see Turner and Laliberté 2015 for a review). We assume that the broad chronology of dune formation and the relative spatial configuration of dunes are consistent along the entire Swan Coastal Plain (Playford *et al.*, 1976;McArthur, 2004). We also assume that the Swan Coastal Plain chronology is broadly comparable to the main stages of dune formation along the Warren chronosequence (Playford *et al.*, 1976), given that the formation of the main coastal dunes in the region has been driven by the same sea-level high stands during interglacial periods throughout the Pleistocene (Kendrick *et al.*, 1991). Thus, time is constrained only within the limits of our broad estimates about timing of dune formation.

### Climate along the chronosequences

Regional variation in precipitation is mapped in Figure 1, while climate data for the four chronosequences are shown in Table 1 and Figure 1. Mean annual rainfall increases from 533 mm at Jurien Bay in the north to 1185 mm at Warren in the south. The dry season, defined as the number of months receiving < 30 mm of rainfall, varies from two months at Warren Beach to seven months at Jurien Bay. Mean annual temperature varies from 15.2°C at Warren in the south to 19.0°C at Jurien Bay in the north. Mean monthly minimum temperatures (January/February) are 10.1°C at Warren and 13.1°C at Jurien Bay, while mean monthly maximum temperatures (July) are 20.3°C at Warren and 25.6°C at Guilderton. The calculated annual potential evapotranspiration ranges from 1133 mm at Warren Beach to 1433 mm at Jurien Bay, reflected in water balances ranging from – 900 mm at Jurien Bay to + 52 mm at Warren.

**Table 1.**
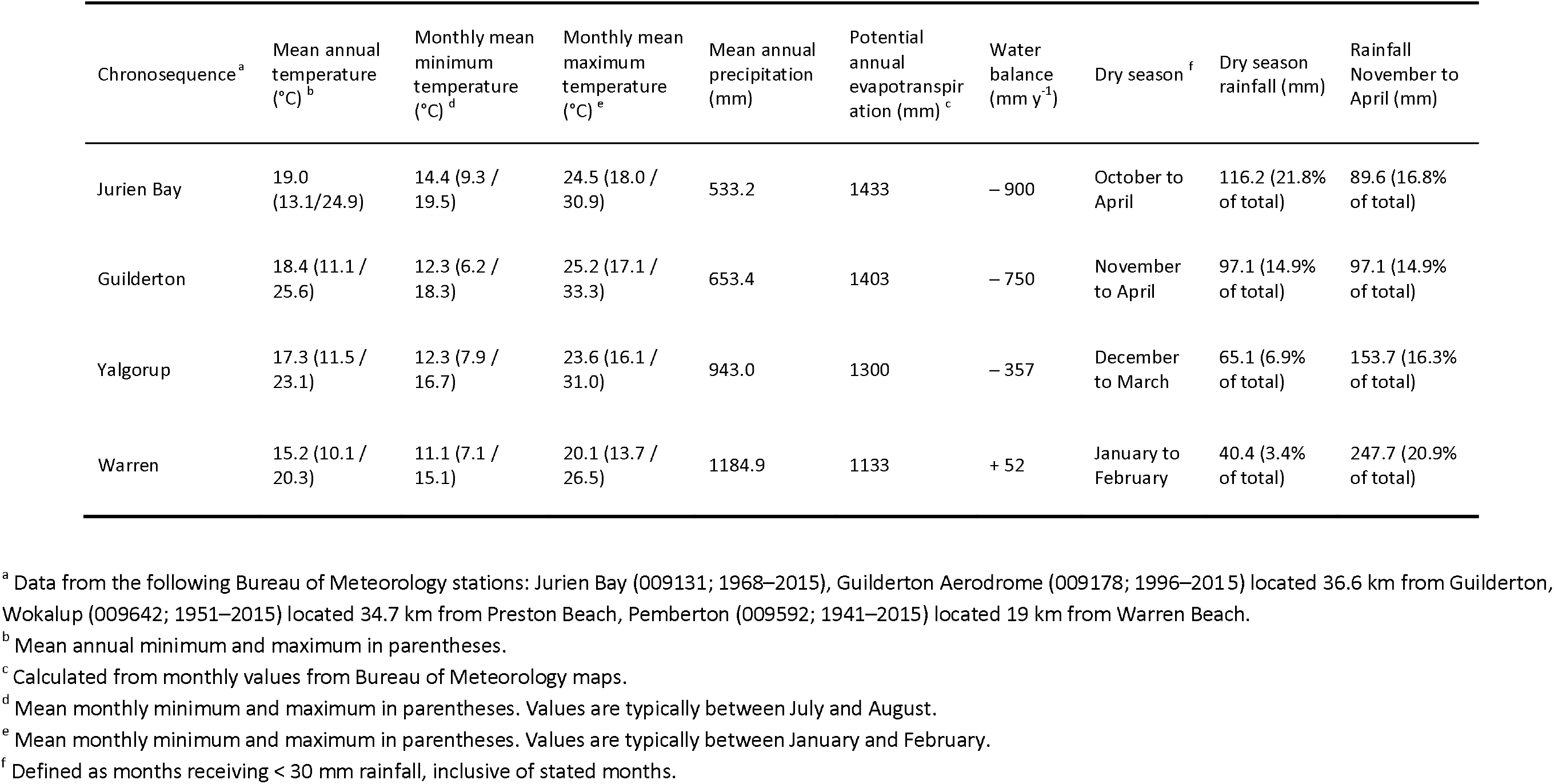
Climate at four long-term chronosequences in southwestern Australia.

| Chronosequence<sup>a</sup> | Mean annual temperature (°C)<sup>b</sup> | Monthly mean minimum temperature (°C)<sup>d</sup> | Monthly mean maximum temperature (°C)<sup>e</sup> | Mean annual precipitation (mm) | Potential annual evapotranspiration (mm)<sup>c</sup> | Water balance (mm y<sup>-1</sup>) | Dry season<sup>f</sup> | Dry season rainfall (mm) | Rainfall November to April (mm) |
| --- | --- | --- | --- | --- | --- | --- | --- | --- | --- |
| Jurien Bay | 19.0 (13.1/24.9) | 14.4 (9.3 / 19.5) | 24.5 (18.0 / 30.9) | 533.2 | 1433 | – 900 | October to April | 116.2 (21.8% of total) | 89.6 (16.8% of total) |
| Guilderton | 18.4 (11.1 / 25.6) | 12.3 (6.2 / 18.3) | 25.2 (17.1 / 33.3) | 653.4 | 1403 | – 750 | November to April | 97.1 (14.9% of total) | 97.1 (14.9% of total) |
| Yalgorup | 17.3 (11.5 / 23.1) | 12.3 (7.9 / 16.7) | 23.6 (16.1 / 31.0) | 943.0 | 1300 | – 357 | December to March | 65.1 (6.9% of total) | 153.7 (16.3% of total) |
| Warren | 15.2 (10.1 / 20.3) | 11.1 (7.1 / 15.1) | 20.1 (13.7 / 26.5) | 1184.9 | 1133 | + 52 | January to February | 40.4 (3.4% of total) | 247.7 (20.9% of total) |

<sup>a</sup> Data from the following Bureau of Meteorology stations: Jurien Bay (009131; 1968–2015), Guilderton Aerodrome (009178; 1996–2015) located 36.6 km from Guilderton, Wokalup (009642; 1951–2015) located 34.7 km from Preston Beach, Pemberton (009592; 1941–2015) located 19 km from Warren Beach.
<sup>b</sup> Mean annual minimum and maximum in parentheses.
<sup>c</sup> Calculated from monthly values from Bureau of Meteorology maps.
<sup>d</sup> Mean monthly minimum and maximum in parentheses. Values are typically between July and August.
<sup>e</sup> Mean monthly minimum and maximum in parentheses. Values are typically between January and February.
<sup>f</sup> Defined as months receiving < 30 mm rainfall, inclusive of stated months.

There is little information on paleoclimate for the four sequences, although it has been argued that the Swan Coastal Plain has been climatically buffered over the lifespan of the sequences (Wyrwoll *et al,* 2014). In particular, Karri trees *(Eucalyptus diversicolor)* and other moisture sensitive plants in the far southwest of the region did not disappear despite arid conditions further inland.

In Soil Taxonomy, soil moisture and temperature regimes are defined by the control section between 30 and 90 cm in sandy textured soils (Soil Survey Staff, 1999). We do not have soil temperature data for the chronosequences, but can estimate soil temperature regimes from air temperatures at nearby stations. Mean annual air temperature varies between 15.2°C at Pemberton (approximately 18 km from the Warren chronosequence) and 19.0°C at Jurien Bay, and for all sequences the difference between the minimum and maximum mean monthly temperature is > 6°C. The soil temperate regime is therefore Thermic for all four chronosequences. Despite the marked variation in rainfall and potential evapotranspiration along the climate gradient, the four chronosequences all under a xeric moisture regime, because the soil is dry for at least 45 days in summer, wet for at least 45 days in the winter, and moist for more than half the year in total. The latter is marginal at Jurien Bay, where precipitation exceeds evapotranspiration for at least six months of the year. In the eeric moisture regime rainfall is particularly effective for leaching, because it occurs in the winter when potential evapotranspiration is lowest.

### Soil sampling

A profile pit was excavated on each of six (Jurien Bay) or seven (all other chronosequences) stages along each chronosequence (Fig. 2). Profile pits were located on the shoulders or upper slopes of dunes, except for the stage-6 profile where dune morphology was indistinct. Pits were at least 1 m deep, and up to 2 m deep on older dunes. Deeper soils were sampled by augering through the pit floor as necessary. However, deep augering was often constrained by dry and incohesive sand that was not retained in the auger, despite the frequent addition of water to the auger hole and the use of a specially designed sand auger (Dormer Soil Samplers, Murwillumbah South, New South Wales, Australia). Soils were described according to Soil Taxonomy (Soil Survey Staff, 1999) and the Australian Soil Classification System (Isbell, 2002) and samples taken from each horizon for bulk density and laboratory analysis. Profile descriptions and analytical data are presented in full in Supplementary Online Material.

To quantify changes in soil nutrients relevant to potential limitation of biological activity, we sampled surface soils (0–10 cm depth) in eighty 10 m × 10 m plots across the four chronosequences *(i.e.* 20 plots per chronosequence). In each chronosequence, we first selected five chronosequence stages (stages 1, 2, 3, 4 and 6) that represented a strong gradient of soil nutrient availability and of the type and strength of nutrient limitation, based on previous work conducted along the Jurien Bay chronosequence (Laliberté *et al.*, 2012;Hayes *et al.*, 2014; Turner & Laliberté, 2015) and on previous soil analyses by McArthur (2004). We excluded the older Spearwood dunes (stages 5a and 5b) from the surface soil sampling as there is relatively little variation in surface soil chemistry between older Spearwood and Bassendean soils (Turner & Laliberté, 2015). For the Jurien Bay chronosequence, we randomly selected four existing plots used in previous studies (Laliberté *et al.*, 2012;Hayes *et al.*, 2014;Laliberté *et al.*, 2014; Turner & Laliberté, 2015;Zemunik *et al.*, 2015). For the other three chronosequences, we positioned four replicate sampling plots in each of the five chronosequence stages at random positions near the profile pits, ensuring that replicate plots followed the same dune. Replicate plots within each chronosequence stage were >50 m apart. In each plot, we collected four soil samples to 20 cm depth using a 50-mm diameter sand auger. Those four soil samples were bulked and homogenised at the plot level prior to chemical analyses. The homogenised samples were sieved (<2 mm) to remove roots and other large organic debris and then air-dried prior to laboratory analyses.

### Soil analysis

Soil analysis for profile pits and surface samples was identical to that described previously (Turner & Laliberté, 2015). Briefly, soil pH was determined in both deionised water and 10 mM CaCl_2_ in a 1:2 soil to solution ratio using a glass electrode. The concentrations of sand (53 μm–2 mm), silt (2 μm– 53 μm), and clay (< 2 μm) sized particles were determined by the pipette method following pretreatment to remove soluble salts and organic matter (Gee & Or, 2002), with further separation of sand fractions by manual dry sieving. Total carbon (C) and N were determined by automated combustion and gas chromatography with thermal conductivity detection using a Thermo Flash 1112 elemental analyser (CE Elantech, Lakewood, NJ). Total P was determined by ignition (550°C, 1 h) and extraction in 1 M H_2_SO_4_ (16 h, 1:50 soil to solution ratio) (Walker & Adams, 1958). Exchangeable cations were determined by extraction in 0.1 M BaCl_2_ (2 h, 1:30 soil to solution ratio), with detection by inductively-coupled plasma optical-emission spectrometry (ICP-OES) on an Optima 7300 DV (Perkin-Elmer Ltd, Shelton, CT) (Hendershot *et al.*, 2008). Carbonate was determined by mass loss after addition of 3 M HCl (Loeppert & Suarez, 1996) and organic C was calculated as the difference between total C and CaCO_3_–C. Bulk density was determined by taking three replicate cores of known volume per horizon using a 7.5 cm diameter stainless steel ring and determining the soil mass after drying at 105°C. Readily-exchangeable phosphate was determined by extraction with anion exchange membranes (resin P) (Turner & Romero, 2009). Total exchangeable bases (TEB) was calculated as the sum of Ca, K, Mg, and Na; effective cation exchange capacity (ECEC) was calculated as the sum of Al, Ca, Fe, K, Mg, Mn, and Na; base saturation was calculated by (TEB ÷ ECEC) × 100.

### Leaf area index

Leaf area index (LAI) was estimated in the same plots from which surface soils were collected using a portable plant canopy imager (Cl-110, CID Bio-Science, Camas, WA, USA). We took four canopy images per plot, each separated by 7 m. Images were taken with the camera as close to possible to the ground surface to include low-stature vegetation and processed using the built-in software. Leaf area index was calculated using the gap-fraction inversion procedure.

### Statistical analysis

Differences in soil properties among chronosequences and chronosequence stages were tested using generalised least squares models using the ‘nlme’ package in R (Pinheiro & Bates, 2000). Chronosequence and chronosequence stages, and the interaction between these two factors, were treated as fixed factors in the models. Model assumptions were assessed visually and appropriate variance structures were specified in the models if they improved the fit, as determined by likelihood-ratio tests; in general, different variances for each chronosequence and stage combination were used. We calculated 95% confidence intervals from the generalised least square models and used those in graphs; the confidence intervals can be used to visually assess statistically significant differences among means in pairwise comparisons if intervals do not overlap.

## Results

### Vegetation

Vegetation varies markedly along the climosequence, from low stature shrubland in the north to relatively tall eucalyptus forest in the south (Fig 3A-D). This change in vegetation structure was reflected in variation in leaf area index, for which maximum values increased from approximately 0.5 or less at Jurien Bay in the drier north to 1.5 at Warren Beach in the wetter south (Fig. 3E). Within the sequences, leaf area index generally increased in younger stages and declined in the older stages, although this was less clear for the relatively dry Jurien Bay chronosequence and most pronounced for the wet Warren Beach chronosequence.

**Figure 3.**
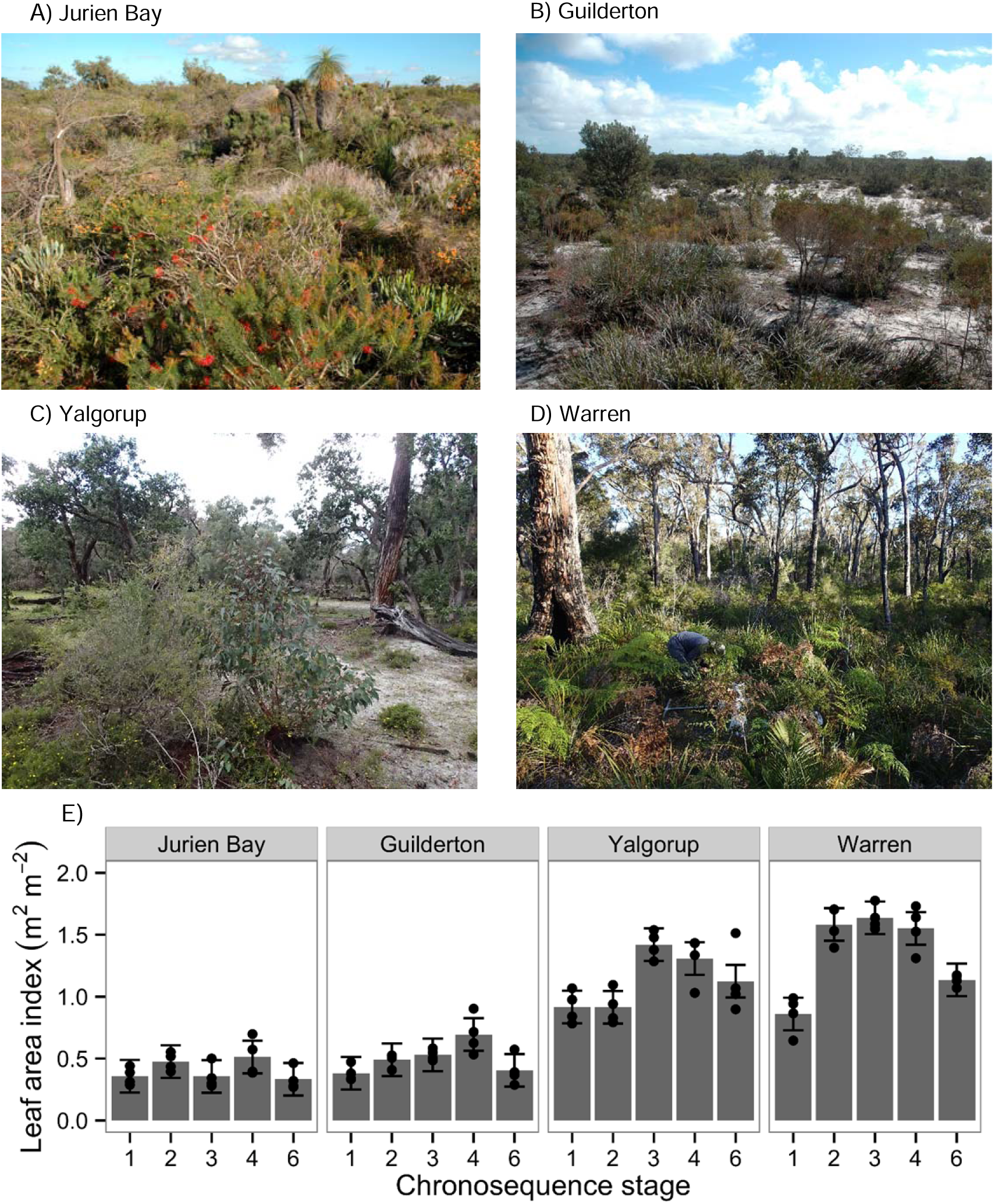
(A-D) Changes in vegetation in the oldest chronosequence stage (i.e. stage 6) for each of the four chronosequences, and (E) leaf area index (LAI) along the four dune chronosequences. Leaf area index was not measured on chronosequence stages 5a and 5b. Bar heights represent means (*n* = 4) and error bars represent 95% confidence intervals from generalised least-square models accounting for different variances among chronosequence stages. Means whose error bars do not overlap can be considered significantly different. Actual values are shown as points. Photo credits: P. Kardol, E. Laliberté, F. Teste, B. Turner and G. Zemunik.

### Pedogenesis along the chronosequences

Pedogenesis along all chronosequences followed a similar pattern to that reported previously for the Jurien Bay chronosequence (Turner & Laliberté, 2015). Young soils developed on Holocene dunes were greyish, yellowish, or pale brown, with weakly developed surface horizons enriched with organic matter overlying several meters of unweathered calcareous sand (see Supplementary Material). All young soils contained carbonate and were strongly or very strongly alkaline. Carbonate concentrations in the beach sand and youngest soils were greatest at Jurien (≥ 80% CaCO_3_), less at Guilderton and Yalgorup (> 40%) and least at Warren (≤5%) (Table 2). This indicates a gradient of declining carbonate concentration in modern parent sand from north to south. Despite this, young soils were of similar strongly alkaline pH throughout the climosequence (Table 2). All soils across the four chronosequences contained > 90% sand (Table 2), predominantly fine, medium, or coarse (0.1 – 1.0 mm), with few very fine or very coarse grains (Table S1). Carbonate concentrations declined with pedogenesis in all four chronosequences, and was completely absent from stage 4 profiles (Middle Pleistocene) above the indurated petrocalcic horizons (calcrete). Carbonate depletion occurred earlier in the surface horizons of the two wetter sequences, being evident in stage 2 profiles at Yalgorup and Warren (Table 2, Fig. S1), and was almost completely leached from the stage 3 (old Holocene) dunes at Warren.

**Table 2.** Chronosequence stages, taxonomic classes and features, and profile-weighted soil physical and chemical properties for soils along the Jurien Bay, Guilderton, Yalgorup, and Warren chronosequences, southwestern Australia. ECEC, effective cation exchange capacity.

| Chronosequence stage | Soil Taxonomy | Carbonate (%) | Sand <sup>a</sup> (%) | Silt (%) | Clay (%) | pH (H<sub>2</sub>O) | pH (CaCl<sub>2</sub>) | ECEC (cmol<sub>c</sub> kg<sup>-1</sup>) | Base sat. (%) |
| --- | --- | --- | --- | --- | --- | --- | --- | --- | --- |
| **JURIEN BAY <sup>b</sup>** |  |  |  |  |  |  |  |  |  |
| 1 | Carbonatic, thermic, Typic Xeropsamments | 82 | 98.2 | 1.1 | 0.7 | 9.1 | 8.2 | 12.93 | 100.0 |
| 2 | Carbonatic, thermic, Typic Xeropsamments | 66 | 95.9 | 2.1 | 2.0 | 9.2 | 8.1 | 5.52 | 99.9 |
| 3 | Siliceous, thermic, Typic Xeropsamments | 25 | 96.9 | 1.4 | 1.6 | 8.9 | 8.0 | 5.01 | 99.4 |
| 4 | Sandy, siliceous, thermic, Calcic Haploxerepts | 0 | 94.0 | 2.9 | 3.1 | 6.9 | 6.1 | 1.94 | 100.0 |
| 5 | Thermic, Xeric Quartzipsamments | 0 | 96.7 | 1.3 | 2.0 | 6.8 | 5.7 | 0.96 | 99.9 |
| 6 | Thermic, Xeric Quartzipsamments | 0 | 97.3 | 2.3 | 0.3 | 5.6 | 4.4 | 0.98 | 97.4 |
| **GUILDERTON** |  |  |  |  |  |  |  |  |  |
| 1 | Carbonatic, thermic, Typic Xeropsamments | 45 | 98.6 | 0.1 | 1.3 | 9.0 | 7.5 | 15.12 | 100.0 |
| 2 | Carbonatic, thermic, Typic Xeropsamments | 49 | 97.7 | 0.7 | 1.5 | 9.1 | 7.7 | 3.04 | 100.0 |
| 3 | Carbonatic, thermic, Typic Xeropsamments | 38 | 92.3 | 2.3 | 5.4 | 8.7 | 7.3 | 5.15 | 100.0 |
| 4 | Thermic, uncoated, Xeric Quartzipsamments | 0 | 93.5 | 2.3 | 4.2 | 6.0 | 5.2 | 0.43 | 98.6 |
| 5a | Thermic, uncoated, Xeric Quartzipsamments | 0 | 95.1 | 1.5 | 3.4 | 5.7 | 4.8 | 0.19 | 87.3 |
| 5b | Thermic, uncoated Xeric Quartzipsamments | 0 | 98.5 | 0.7 | 0.9 | 5.5 | 4.7 | 0.11 | 85.6 |
| 6 | Thermic, uncoated, Xeric Quartzipsamments | 0 | 98.6 | 0.6 | 0.8 | 5.6 | 3.8 | 0.68 | 100.0 |
| **YALGORUP** |  |  |  |  |  |  |  |  |  |
| 1 | Mixed, thermic, Typic Xeropsamments | 37 | 98.4 | 0.6 | 1.0 | 8.6 | 6.9 | 3.08 | 100.0 |
| 2 | Carbonatic, thermic, Typic Xeropsamments | 48 | 96.8 | 0.4 | 2.8 | 8.6 | 7.5 | 3.98 | 99.8 |
| 3 | Carbonatic, thermic, Typic Xeropsamments | 42 | 94.0 | 1.5 | 4.5 | 8.9 | 7.6 | 7.21 | 99.9 |
| 4 | Sandy, siliceous, thermic, Calcic Haploxerepts | 0 | 97.4 | 1.2 | 1.3 | 6.8 | 6.3 | 1.13 | 98.7 |
| 5a | Thermic, uncoated, Xeric Quartzipsamments | 0 | 96.5 | 2.0 | 1.5 | 6.2 | 5.0 | 0.44 | 91.5 |
| 5b | Thermic, uncoated, Xeric Quartzipsamments | 0 | 98.0 | 1.1 | 0.9 | 5.9 | 4.2 | 0.16 | 84.0 |
| 6 | Thermic, uncoated, Xeric Quartzipsamments | 0 | 97.1 | 2.4 | 0.5 | 5.9 | 4.1 | 0.30 | 94.3 |
| **WARREN** |  |  |  |  |  |  |  |  |  |
| 1 | Thermic, uncoated, Xeric Quartzipsamments | 4.2 | 99.1 | 0.5 | 0.4 | 9.2 | 8.1 | 3.79 | 99.9 |
| 2 | Thermic, uncoated, Xeric Quartzipsamments | 1.5 | 99.1 | 0.3 | 0.6 | 9.0 | 8.1 | 2.36 | 100.0 |
| 3 | Thermic, uncoated, Xeric Quartzipsamments | 0.2 | 97.8 | 1.7 | 0.5 | 6.2 | 5.3 | 1.05 | 93.5 |
| 4 | Thermic, uncoated, Xeric Quartzipsamments | 0 | 98.9 | 0.6 | 0.5 | 5.8 | 4.9 | 0.35 | 80.1 |
| 5a | Thermic, uncoated, Xeric Quartzipsamments | 0 | 98.9 | 0.6 | 0.6 | 5.9 | 4.7 | 0.51 | 94.4 |
| 5b | Thermic, uncoated, Xeric Quartzipsamments | 0 | 98.9 | 0.6 | 0.5 | 5.3 | 4.2 | 0.68 | 97.2 |
| 6 | Thermic, uncoated, Xeric Quartzipsamments | 0 | 98.8 | 0.7 | 0.5 | 5.3 | 4.0 | 0.27 | 94.9 |

As pedogenesis proceeds and soils begin to acidify the carbonate is progressively leached from the profiles and precipitated at depth, eventually forming an indurated petrocalcic horizon beneath residual yellow sands, characteristic of young Spearwood dunes (i.e. 120,000 years old; stage 4). These soils have brownish yellow subsoil consisting of > 95% residual quartz sand. The yellow colour derives from goethite and the weathering of heavy minerals (Bastian, 1996). Continued pedogenesis leads to progressive leaching of iron oxides, sometimes forming a Bs horizon, and eventually leaving bleached quartz sand profiles several meters deep, characteristic of Bassendean dunes (i.e. approximately 2 million years old; stage 6). These oldest soils have extremely low cation exchange capacity, although base saturation remains high (Table 2).

The major pedogenic transitions were observed only weakly at Jurien Bay, but were captured clearly in profiles at Guilderton and Yalgorup (Fig. 4). In particular, the old Quindalup dunes (stage 3) at Guilderton exhibited a calcic horizon with weakly cemented secondary carbonates and an incipient undulating petrocalcic horizon (Fig. 4c-e, 4j-k). In contrast, pedogenesis at Warren did not involve the formation of a petrocalcic horizon, presumably due to the lower carbonate content of the parent sand and the greater leaching potential. The old Spearwood dunes at Guilderton, Yalgorup, and Warren all included clear examples of the transition from Spearwood to Bassendean soils, with gradual leaching of iron oxides leading to the development of deep bleached eluvial (E) horizons over yellow sand (Fig. 4f-i, 4l-n). Previous studies at Jurien captured only the initial stage of this process, with a shallow incipient eluvial horizon in the oldest Spearwood dune.

**Figure 4.**
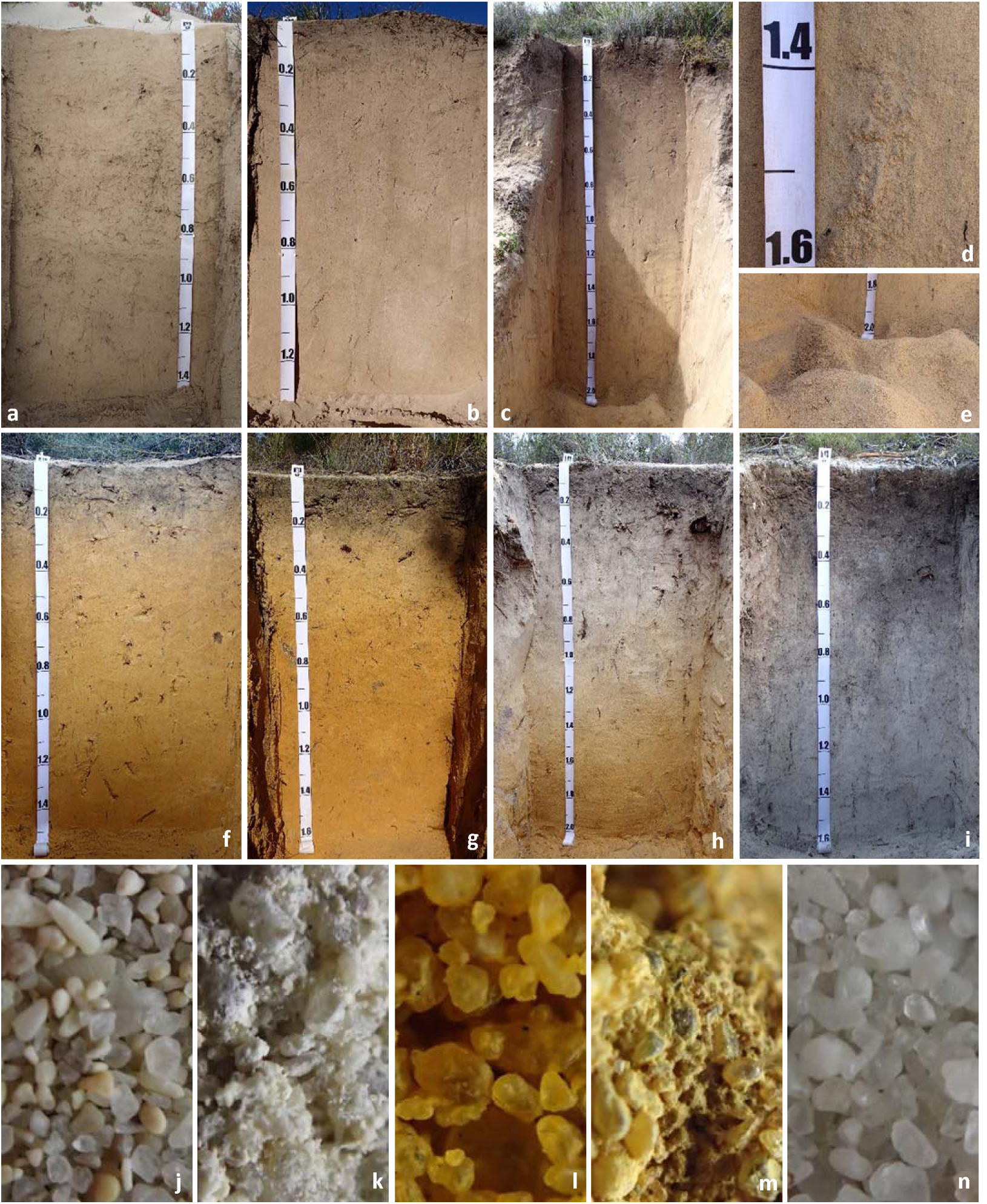
Soil development along the Guilderton chronosequence: (a) young Holocene dune (∼100 years, stage 1), (b) medium-aged Holocene dune with incipient A horizon (∼1000 years, stage 2), (c) old Holocene dune (6500 years, stage 3), (d) precipitation of pedogenic carbonate forming a calcic horizon in old Holocene dune, (e) surface of a petrocalcic horizon in old Holocene dune, (f) young Middle Pleistocene dune showing decalcified sand coated with iron oxides (120,000 years, stage 4), (g) medium-aged Middle Pleistocene dune with incipient E horizon (∼250,000 years, stage 5a), (h) old Middle Pleistocene dune with 1 m deep bleached E horizon (∼400,000 years, stage 5b), (i) Early Pleistocene dune showing bleached quartz sand (∼2 million years, stage 6). Fine detail in subsoil of five profiles showing key pedogenic stages: (j) a mixture of carbonate and quartz sand-sized grains in young Holocene dune, (k) sand grains cemented by pedogenic carbonate at 150 cm in old Holocene dune, (l) iron oxide coatings on quartz grains in young Middle Pleistocene dune, (m) quartz grains cemented by iron oxide at 250 cm in old Middle Pleistocene dune, (n) clean quartz grains leached of iron oxides in Early Pleistocene dune.

### Soil Taxonomy

Profiles along all four sequences did not generally qualify as Inceptisols because the particle size classes were too coarse for cambic horizons (which require a texture of very fine sand or finer; Table 2). As at Jurien Bay, young soils (stages 1–3; Fig. 4a-c) along all four sequences were classified as Psamments (Entisols) because of their coarse sandy texture. Due to the nature of the sand, these young profiles were Quartzipsamments at Warren, but Xeropsamments elsewhere (Table 2). The Xeropsamments were carbonatic, except the youngest soil at Yalgorup, which contained marginally insufficient carbonate in the profile. The oldest Holocene soils at Guilderton and Yalgorup did not qualify as Inceptisols, because the calcic horizons with free secondary carbonates were not within 100 cm of the soil surface. In the Australian Soil Classification System (Isbell 2002), the young Holocene soils are predominantly Shelly or Arenic Rudosols, while the old Holocene soils are Calcic or Petrocalcic Tenosols at Guilderton and Yalgorup, and Grey-Orthic Tenosols at Warren.

Middle and Early Pleistocene soils (stages 4–6; Fig. 4f-i) were Quartzipsamments (Entisols) along all four sequences due to their deep quartz sand profiles lacking diagnostic horizons in the upper part of the profiles. The exceptions were the youngest Spearwood soils at Jurien and Yalgorup, which qualified as Calcic Haploxerepts (Inceptisols) because of the petrocalcic within 150 cm of the soil surface. The stage 4 profile at Guilderton was similar, with contained a petrocalcic horizon below yellow sand, but did not qualify as an Inceptisol because the petrocalcic horizon was > 150 cm below the soil surface. In the Australian Soil Classification system the Middle Pleistocene soils qualify as Grey or Yellow-Orthic Tenosols, with the younger profiles being in the Petrocalcic Great Group, while the older soils with a spodic horizon (stages 5b and 6) qualify as Sesquic or Humosesquic Aerie Podosols.

Soils from stage 6 (Fig 4i) were morphologically, chemically, and physically similar at each of the four sequences. The designation as Entisols (i.e. young soils) for these 2-million year profiles is an artifact of the Soil Taxonomy system, which does not consider parts of the profile deeper than 200 cm (Turner & Laliberté, 2015). We observed a clear spodic horizon in Bassendean soils only at the Yalgorup chronosequence, although this indurated coffee rock was too deep (>200 cm) for this profile to be placed in the Spodosols. An additional profile at Warren, not on the main chronosequence but on Bassendean sand, qualified as a Spodosol due to a spodic horizon within 200 cm of the soil surface (see profile classifications in Supplementary Material). Quartz-rich soils often contain insufficient metals to yield a spodic horizon and therefore develop into Quartzipsamments (Fanning & Fanning, 1989), although variation in the presence of a spodic horizon in the Swan Coastal Plain has also been linked to differences in the proximity to the water table (McArthur and Russell 1978). In the absence of spodic horizons, the Bassendean profiles qualify as Grey-Orthic-Tenesols in the Australian Soil Classification System.

### Organic carbon, total nitrogen, and total phosphorus

Profile-weighted organic C and N stocks followed a consistent pattern along chronosequences, but were variable across the climosequence (Table 3, Fig. S2, Fig. S3). Stocks of both C and N tended to increase initially to greatest amounts in old Holocene dunes, being as high as 12 kg C m^−2^ / 648.5 g N m^−2^ at Yalgorup (Table 3), and then declined to lower amounts on older dunes. However, organic C was noticeably high in the oldest soil at Guilderton (9.8 kg C m^−2^) compared to the earlier stages of the sequence.

**Table 3.** Profile weighted nutrient concentrations for the upper 100 cm of soil in profile pits along the Jurien Bay, Guilderton, Yalgorup, and Warren chronosequences, southwestern Australia. Data for the Jurien Bay chronosequence are from Turner & Laliberté (2015).

| Chronosequence stage | Organic C (g m<sup>-2</sup>) | Total N (g m<sup>-2</sup>) | Total P (g m<sup>-2</sup>) | C:N | C:P | N:P |
| --- | --- | --- | --- | --- | --- | --- |
| **JURIEN BAY** |  |  |  |  |  |  |
| 1 | 8900 | 495.5 | 384.3 | 18.0 | 23.2 | 1.3 |
| 2 | 14678 | 763.3 | 346.4 | 19.2 | 42.4 | 2.2 |
| 3 | 17720 | 362.2 | 194.8 | 48.9 | 90.9 | 1.9 |
| 4 | 3864 | 166.5 | 28.8 | 23.2 | 134.4 | 5.8 |
| 5 | 2917 | 119.2 | 12.7 | 24.5 | 228.9 | 9.4 |
| 6 | 4063 | 117.8 | 6.4 | 34.5 | 639.2 | 18.5 |
| **GUILDERTON** |  |  |  |  |  |  |
| 1 | 2165 | 223.3 | 240.1 | 9.7 | 9.0 | 0.9 |
| 2 | 4284 | 371.4 | 136.3 | 11.5 | 31.4 | 2.7 |
| 3 | 7282 | 537.7 | 198.5 | 13.5 | 36.7 | 2.7 |
| 4 | 1500 | 110.8 | 14.5 | 13.5 | 103.3 | 7.1 |
| 5a | 1924 | 102.0 | 10.9 | 18.9 | 177.2 | 9.4 |
| 5b | 1772 | 54.3 | 3.1 | 32.6 | 565.6 | 17.3 |
| 6 | 9801 | 190.7 | 2.6 | 51.4 | 3764.4 | 73.2 |
| **YALGORUP** |  |  |  |  |  |  |
| 1 | 3324 | 224.5 | 210.0 | 14.8 | 15.8 | 1.1 |
| 2 | 6030 | 503.0 | 219.5 | 12.0 | 27.5 | 2.3 |
| 3 | 12162 | 648.6 | 277.7 | 18.7 | 43.8 | 2.3 |
| 4 | 4922 | 200.8 | 31.4 | 24.5 | 156.9 | 6.4 |
| 5a | 3336 | 117.5 | 6.6 | 28.4 | 508.7 | 17.9 |
| 5b | 6153 | 151.8 | 4.3 | 40.5 | 1434.8 | 35.4 |
| 6 | 5430 | 181.0 | 5.6 | 30.0 | 976.7 | 32.6 |
| **WARREN** |  |  |  |  |  |  |
| 1 | 188 | 47.9 | 35.6 | 3.9 | 5.3 | 1.3 |
| 2 | 2055 | 145.7 | 23.0 | 14.1 | 89.2 | 6.3 |
| 3 | 3337 | 239.5 | 18.4 | 13.9 | 181.4 | 13.0 |
| 4 | 1925 | 99.4 | 11.4 | 19.4 | 169.5 | 8.8 |
| 5a | 2820 | 148.3 | 4.4 | 19.0 | 636.3 | 33.5 |
| 5b | 3932 | 140.2 | 4.5 | 28.0 | 869.0 | 31.0 |
| 6 | 2675 | 74.1 | 2.3 | 36.1 | 1141.1 | 31.6 |

Total P stocks in the youngest soils (stage 1) were greatest at Jurien (300 g P m-2), lower at Guilderton and Yalgorup (210–240 g P m^−2^), and lowest at Warren (36 g P m^−2^), reflecting the carbonate and P content of the parent sand (Table 3, Fig. 5). With the exception of an increasing total P stock in Holocene dunes at Yalgorup, presumably due to high productivity at the old Holocene site (reflected in the high organic C stock), total P declined more or less continuously throughout the chronosequences, reaching extremely low amounts (< 5 g P m^−2^) on the oldest soils Spearwood and Bassendean soils.

**Figure 5.**
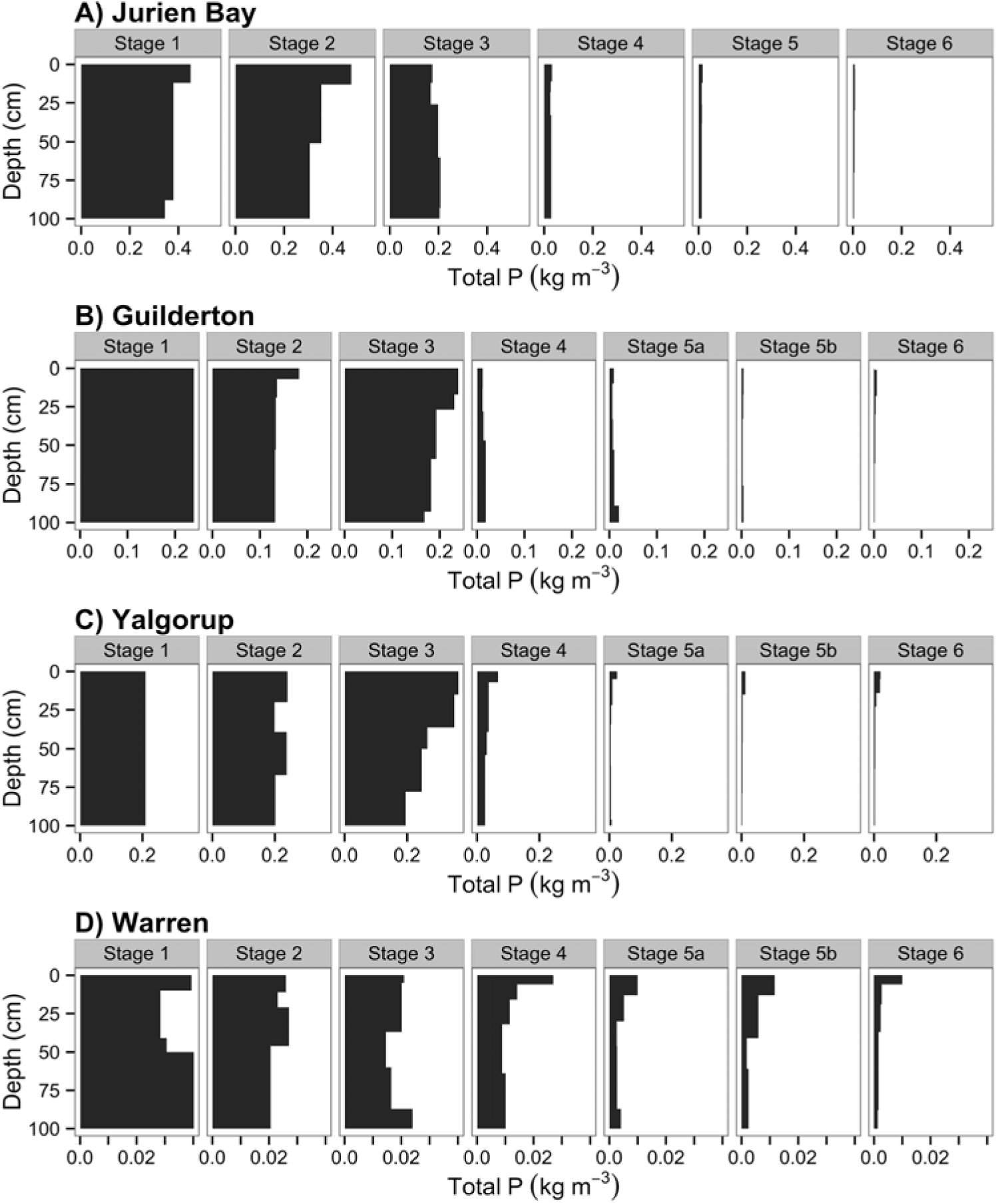
Profile-weighted contents of total P to 100 cm depth along the four dune chronosequences: (A) Jurien Bay, (B) Guilderton, (C) Yalgorup and (D) Warren. Note the differences in the x-axis values among chronosequences.

In surface soils (0–10 cm depth) along all four sequences, organic C in surface soils was lowest in the youngest chronosequence stage and then increased to reach maximal values in stage 2 or 3, except for Yalgorup where organic C was also high in the oldest Holocene stage (Fig. 6A). There was a decline in organic C from stage 3 to 4 in three chronosequences (Jurien Bay, Guilderton and Yalgorup), while organic C remained constant from stage 3 to 4 in Warren (Fig. 6A). Total nitrogen followed similar patterns to organic C (Fig. 6B). Total N started at low concentrations in all four chronosequences, then reached maximal values in stage 2 (Jurien Bay) or stage 3 (all other chronosequences). Following this increase, there was a decline in total nitrogen from stage 3 or 4 in all chronosequences (Fig. 6B). In the oldest soils (stage 6), total N reached levels as low as stage 1 for all sequences except Yalgorup (Fig. 6B).

**Figure 6.**
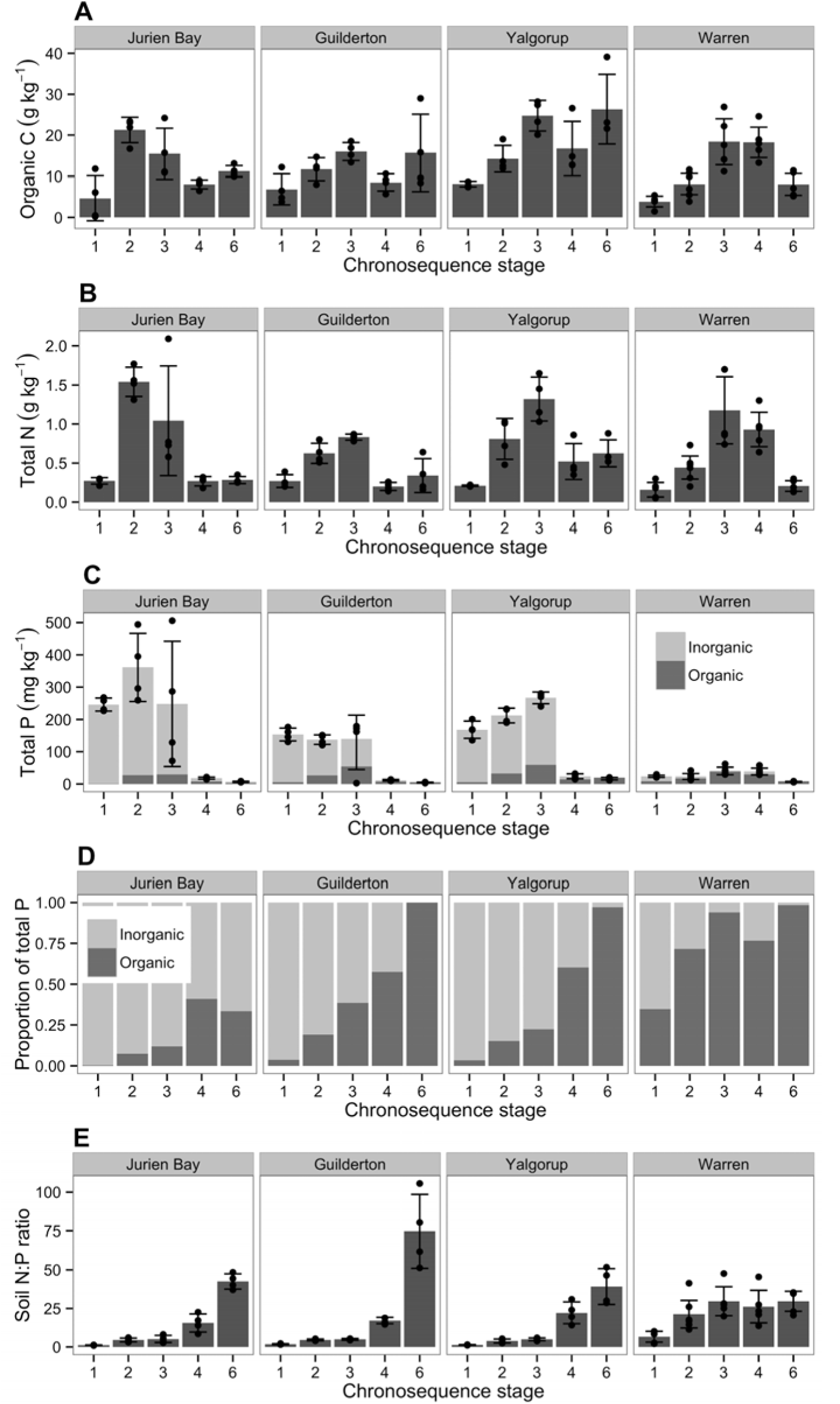
Changes in (A) organic C, (B) total N, (C) total P, (D) proportion of total P present as organic P, and (E) total N: total P ratio, in surface soils (0–10 cm depth) across the four chronosequences. Bar heights represent means (*n* = 4) and error bars represent 95% confidence intervals from generalised least-square models accounting for different variances among chronosequence stages. Means whose error bars do not overlap can be considered significantly different. Actual values are shown as points.

In the three sequences from the Swan Coastal Plain (Jurien Bay, Guilderton, Yalgorup), there were large differences in total P concentrations between the Holocene dunes, which had relatively high total P concentrations, and the older stages, which had extremely low total P concentrations (Fig. 6C). By contrast, total P concentrations in surface soils at Warren were much lower initially (i.e. in stage 1) and increased slightly until stage 4, presumably due to plant uplift and concentration in organic matter at the surface (Laliberté *et al.*, 2012; Turner & Laliberté, 2015), but declined to extremely low concentrations in stage 6 (Fig. 6C).

Along all four chronosequence stages there was a clear increase in the proportion of total P present as organic P during pedogenesis, with organic P reaching almost 100% of surface soil total P in stage 6 for the three wetter sequences (Guilderton, Yalgorup and Warren; Fig. 6D). In general, organic P in surface soils represented a greater proportion of total P as the climate became cooler and wetter (i.e from Jurien Bay to Warren; Fig. 6D).

There were marked increases in surface soil N:P ratio from youngest to oldest stage in all chronosequences (Fig. 6E), eventually reaching high values in the oldest stages (i.e. around 50 or more). The increase was more pronounced in the three drier Swan Coastal Plain chronosequences, whereas N:P increased from stage 1 to 2 at Warren but then did not increase further in older soils (Fig. 6E). Profile-weighted nutrient ratios demonstrated a similar pattern in surface soils (Table 3; Fig. S4). Overall, C:N, C:P, and N:P ratios increased continuously along all four chronosequences, reaching highest values on the oldest soils. For profile-weighted ratios, the greatest values were a C:N ratio of 51.4, C:P ratio of 3764, and N:P ratio of 73.2 in the oldest profile at Guilderton.

### Surface soil pH, exchangeable phosphorus, and exchangeable cations

Surface soils in all four chronosequence stages were initially alkaline (pH in 0.01 M CaCl_2_ > 7) but then gradually acidified during pedogenesis (Fig. 7A). Soil pH in surface soils in stage 1 decreased from the driest to the wettest sequence (i.e. Jurien Bay > Guilderton > Yalgorup > Warren), reflecting the decline in carbonate concentration of the beach sand from Jurien Bay to Warren (Table 2). Similarly, soil pH in the oldest stages was highest at Jurien compared to the other sequences.

**Figure 7.**
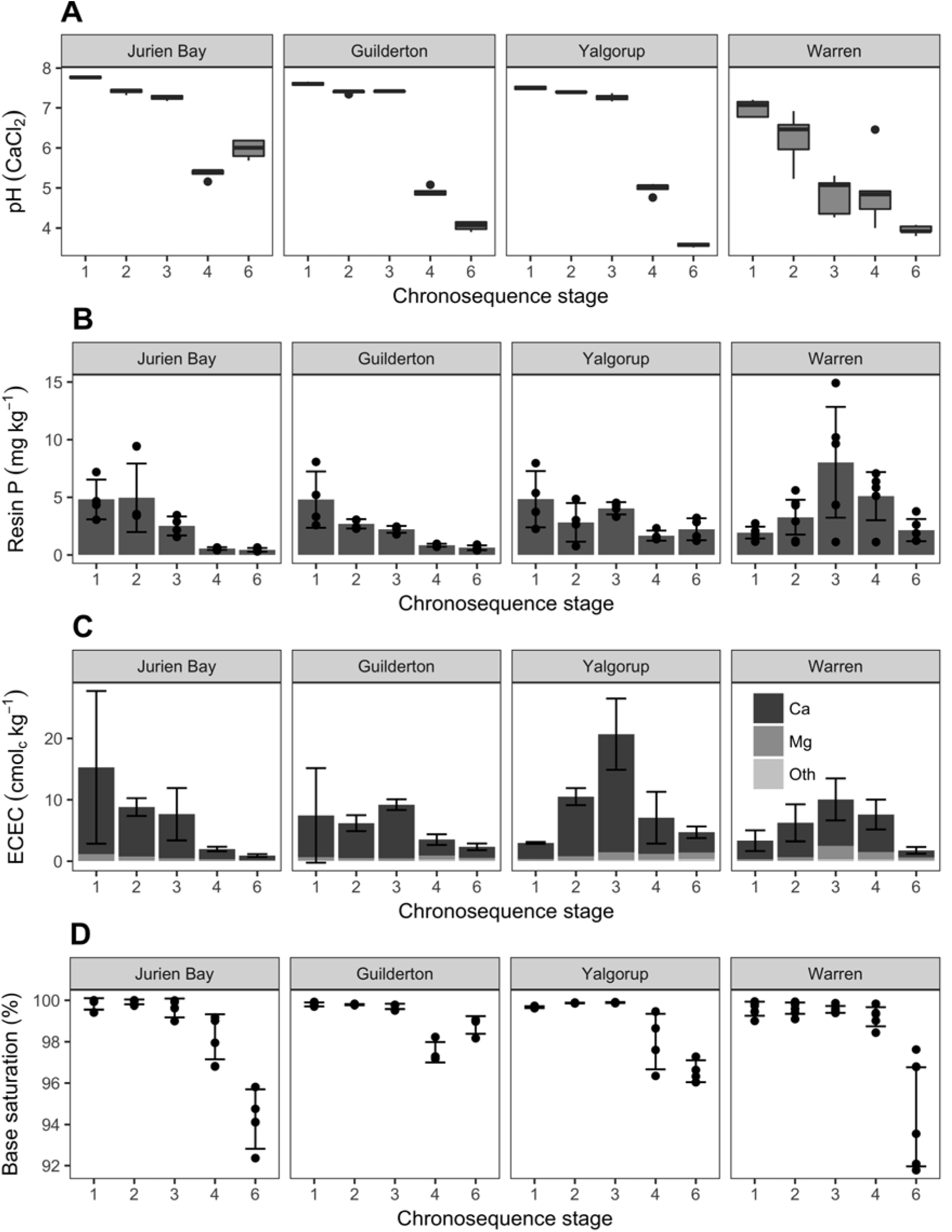
Changes in (A) pH (measured in 0.01 M CaCl_2_), (B) resin-extractable P, (C) exchangeable cations and effective cation exchange capacity (ECEC), and (D) base saturation, in surface soils (0–10 cm depth) across the four chronosequences. In (A), the central horizontal bar in each box shows the median, the box represents the interquartile range, the whiskers show the location of the most extreme data points that are still within a factor of 1.5 of the upper or lower quartiles, and the black points are values that fall outside the whiskers. In B and C, bar heights represent means (*n* = 4) and error bars represent 95% confidence intervals from generalised least-square models accounting for different variances among chronosequence stages. Means whose error bars do not overlap can be considered significantly different. In B, actual values are shown as points.

Exchangeable phosphorus determined by extraction with anion-exchange resins was highest in the youngest soils of the three drier chronosequences and then declined continuously with pedogenesis, reaching lowest values on the oldest soils (Fig. 7B). In contrast, resin phosphate concentrations at Warren increased initially in Holocene dunes, then declined on Middle and Early Pleistocene dunes (Fig. 7B). Overall, resin P concentrations on the oldest dunes were lower at the driest site (Jurien Bay).

In all four chronosequences, effective cation exchange capacity declined in the oldest soils to reach very low levels (Fig. 7C). However, changes in effective cation exchange capacity during the first three stages differed among chronosequences. In Jurien Bay, and in Guilderton to a lesser degree, there was a general decline in effective cation capacity from the youngest to the oldest soil (Fig. 7C). By contrast, in Yalgorup and Warren, there were increases in effective cation exchange capacity from stages 1 to 3 prior to the decline in the older soils (Fig. 7C). In all stages of all four chronosequences, effective cation exchange capacity was largely dominated by Ca irrespective of soil age, while Mg was the second most important exchangeable cation (Fig. 7C). By contrast, other exchangeable cations (K, Fe, Mn, Na, and Al) were present in very small amounts (Fig. 7C). Base saturation was > 90% in surface soils throughout the four sequences, only being < 100% in Pleistocene soils (Fig. 7D).

## Discussion

We define a series of four long-term chronosequences that form a climosequence of chronosequences in southwestern Australia. The system is unique in terms of its Mediterranean climate and its location in a global biodiversity hotspot (Hopper & Gioia, 2004). These four chronosequences allow examination of the influence of climate on patterns of pedogenesis and nutrient status during long-term ecosystem development. Such climate × age systems are exceedingly rare worldwide – the only comparable framework we are aware of is on the Hawaiian Islands, where steep rainfall gradients, well-constrained parent material, and uniform vegetation allow unparalleled control of soil forming factors (Vitousek, 2004).

Despite relatively large changes in climate, pedogenic change was remarkably consistent along the four chronosequences, with decalcification in Holocene dunes and leaching of iron oxides from Middle Pleistocene dunes, yielding bleached quartz sand profiles several meters deep on Early Pleistocene dunes. These transitions were captured by profiles along the chronosequences, providing evidence that soil variation in the sandplains is a result of in situ soil development. Changes in soil nutrients during ecosystem development were also consistent along the four chronosequences, with increasing N concentrations in young soils, declining concentrations of P and exchangeable cations in old soils, and increasing soil N:P ratios throughout ecosystem development. This pattern of soil nutrients therefore corresponds closely with the Walker and Syers (1976)model of nutrient transformations during pedogenesis. Despite clear effects on vegetation structure (as reflected by changes in LAI), climate effects on soil nutrients appeared marginal, with the only clear variation in soil nutrients with climate manifesting in the nature of the soil P pools. Specifically, the proportion of the soil P present in organic forms was greater, and increased earlier in ecosystem development, in wetter sites, while resin P concentrations were also greater in old Holocene and Middle Pleistocene dunes at the wettest site. These differences suggest that greater plant biomass or productivity maintains P in actively cycling pools, and therefore might buffer against nutrient loss by leaching (Porder & Chadwick, 2009).

Vitousek and Chadwick (2013b) describe ‘soil process domains’, in which soil properties change relatively little despite variation in rainfall. These domains occur between ‘pedogenic thresholds’, where soil properties change rapidly over across a relatively small variation in rainfall. They identified three domains for Andisols about 150,000 years old on Hawaii: (i) a low rainfall domain where evapotranspiration exceeds precipitation and carbonate precipitation dominates; (ii) an intermediate domain where biological uplift of nutrients buffers against loss by leaching, (iii) a high rainfall domain where leaching losses dominate. The intermediate rainfall domain on Hawaii occurs between about 700 and 1700 mm and, based on the results for our Australian climosequence, it appears that the four chronosequences occupy a single process domain: patterns of nutrient availability and pedogenesis are relatively similar across the entire climosequence, and precipitation exceeds potential evapotranspiration only at the most southerly site, and there only marginally (52 mm). The lack of variation in patterns of base cations with climate is probably related in part to the sandy nature of the soils, with limited formation of clays or secondary minerals following decalcification. Soil pH and Ca concentrations decline in young soils associated with the depletion of carbonates, exchangeable Al occurs in trace concentrations, and base saturation is therefore always high despite low concentrations of base cations. Indeed, once carbonates are depleted from Holocene soils, and iron oxide coatings leached from the Middle Pleistocene soils, there is essentially no cation exchange capacity other than on organic matter.

Vegetation structure varied markedly along the climosequence and within chronosequences. Along the climosequence, vegetation changed from low-stature shrubland in the drier north to relatively tall *Eucalyptus* forest in the wetter south. Given that changes in fertility with increasing soil age appear relatively constant across the four chronosequences, the overall increase in biomass and leaf area index with increasing rainfall is presumably related to water availability. Patterns in vegetation along the chronosequences (i.e. during ecosystem development) was consistent with the concept of retrogression (Wardle *et al.*, 2004), with LAI initially increasing in the early stages of pedogenesis, and then declining on the oldest soils under strong P limitation. This trend was clearest along the wetter chronosequences, where maximum LAI also occurred earlier in ecosystem development, and least clear in the driest chronosequence, where there was no consistent trend in LAI along the chronosequence. Studies so far at Jurien Bay indicate that P availability constrains ecosystem processes and favours plant species with high P-use efficiency on the oldest soils (Laliberté *et al.*, 2012;Hayes *et al.*, 2014), consistent with the concept of retrogression (Peltzer *et al.*, 2010), even if LAI varies little across the chronosequence. For example, the oldest soils are dominated by slow-growing plant species with long-lived leaves that maintain extremely low foliar P concentrations and a high resorption efficiency for P (Hayes *et al.*, 2014).

Ecosystem retrogression therefore appears to be expressed more strongly in the vegetation structure along wetter chronosequences. Along the Australian climosequence, this variation in expression of strong P limitation on the plant community might be influenced by corresponding variation in plant community diversity. Although we have so far not conducted detailed vegetation assessments along all four chronosequences (but see Zemunik *et al.*, 2015; 2016), our field observations indicate declining diversity with increasing rainfall, from the hyperdiverse shrubland at Jurien Bay (Laliberté *et al.*, 2014;Zemunik *et al.*, 2016) to the comparatively low diversity forest at Warren Beach (Hopper & Gioia, 2004). It has been proposed that ecosystem retrogression is less likely to occur in diverse plant communities such as lowland tropical forests because there are more likely to be species that can maintain productivity on low P soils (Kitayama, 2005). For example, the response of the vegetation to changes in nutrient status linked to long-term pedogenesis is relatively clear along the Hawaiian Island chronosequence where forests are dominated by a single species of tree (Vitousek & Farrington, 1997). In contrast, strongly weathered soils at Jurien Bay support a diverse array of species that exhibit high P-use efficiency and can therefore maintain relatively high biomass on extremely infertile soils (Hayes *et al.*, 2014;Lambers *et al.*, 2015;Zemunik *et al.*, 2015). High regional plant diversity might therefore buffer against a decline in productivity related to the long-term decline in P availability during retrogression (Vitousek, 2004).

There are two caveats concerning the chronosequences. First, there is variation in the parent sand along the coastline, with lower carbonate concentrations, and therefore total P concentrations, towards the south. This regional variation in the chemical composition of beach sand is presumably related to differences in offshore productivity (McArthur, 2004), although we do not have information on whether the pattern in modern sand composition also occurred historically. Second, although we find consistent patterns of soil development along the four chronosequences, we have not so far been able to precisely quantify dune ages and therefore rates of soil development. However, the relative dune ages are fairly well constrained, particularly for the Swan Coastal Plain (see also Turner & Laliberté, 2015), giving confidence that the overall patterns are consistent among the four chronosequences. In the Spearwood dunes, Bastian (1996)recognised five stages, although at least seven sea level high stands are recognised. Aside from the 120,000 year high stand (Marine Isotope Stage 5e) that is relatively easy to identify (and consistent among our chronosequences), a 200,00 year high stand was relatively small and likely to have been over-ridden by the subsequent 120,000 year event. Before that, high stands occurred at approximately 220,000, 240,000, 280,000, 330,000, and 410,000 years, so it is likely that Spearwood stages 4 and 5a in our sequences are separated by at least 100,000 years, but that stages 5a and 5b could be separated by as little as 20,000 years or as much as 190,000 years.

Despite these limitations, we consider that the series of four long-term retrogressive chronosequences across a clear climate gradient provides an important model system for studying long-term soil and ecosystem development, particularly since well-characterised retrogressive sequences are rare worldwide (Peltzer *et al.*, 2010). These four sequences should be particularly valuable as strong natural gradients of soil fertility to explore plant adaptations to declining nutrient availability (e.g.Hayes *et al.*, 2014;Zemunik *et al.*, 2015) and edaphic drivers of plant (Laliberté *et al.*, 2014;Teste *et al.*, 2017) and microbial diversity (Krüger *et al.*, 2015;Albornoz *et al.*, 2016) under contrasting climates, and to examine the independent influence of climate on above and below ground organisms while controlling for variation in soil nutrients.

## Acknowledgements

Funding was provided by a Discovery Early Career Researcher Award (DE120100352) and a Discovery Project (DP130100016) from the Australian Research Council, a Research Collaboration Award from The University of Western Australia to EL and BLT, and a NSERC Discovery Grant awarded to EL. The authors thank Dayana Agudo, Pedro Araúz, Aleksandra Bielnicka, and Paola Escobar for laboratory support, and Felipe Albornoz, Hans Lambers, Kenny Png, François Teste, Karl-Heinz Wyrwoll, and Graham Zemunik for assistance in the field. Figure 2 was created using base maps provided by the Department of Agriculture and Food of Western Australia.

## References

Albornoz, F.E., Teste, F.P., Lambers, H., Bunce, M., Murray, D.C., White, N.E. & Laliberté, E. 2016. Changes in ectomycorrhizal fungal community composition and declining diversity along a 2- million-year soil chronosequence. Molecular Ecology, 25, 4919–4929.

Bastian, L.V. 1996. Residual soil mineralogy and dune subdivision, Swan Coastal Plain, Western Australia. Australian Journal of Earth Sciences, 43, 31–44.

Chadwick, O.A. & Chorover, J. 2001. The chemistry of pedogenic thresholds. Geoderma, 100, 321–353.

Chadwick, O.A., Gavenda, R.T., Kelly, E.F., Ziegler, K., Olson, C.G., Elliott, W.C. & Hendricks, D.M. 2003. The impact of climate on the biogeochemical functioning of volcanic soils. Chemical Geology, 202, 195–223.

Fanning, D.S. & Fanning, M.C.B. 1989. Soil Morphology, Genesis and Classification. John Wiley & Sons, New York.

Feng, J., Turner, B.L., Lü, X., Chen, Z., Wei, K., Tian, J., Wang, C., Luo, W. & Chen, L. 2016. Phosphorus transformations along a large-scale climosequence in arid and semiarid grasslands of northern China. Global Biogeochemical Cycles, 30, 1264–1275.

Gee, G.W. & Or, D. 2002. Particle size analysis. In. Methods of Soil Analysis, Part 4 – Physical Methods (eds. Dane, J.H. & Topp, C.), pp. 255–293. Soil Science Society of America, Madison, WI.

Hayes, P., Turner, B.L., Lambers, H. & Laliberté, E. 2014. Foliar nutrient concentrations and resorption efficiency in plants of contrasting nutrient-acquisition strategies along a 2-million-year dune chronosequence. Journal of Ecology, 102, 396–410.

Hendershot, W.H., Lalande, H. & Duquette, M. 2008. Chapter 18. Ion exchange and exchangeable cations. In. Soil Sampling and Methods of Analysis (eds. Carter, M.R. & Gregorich, E.), pp. 173–178. Canadian Society of Soil Science and CRC Press, Boca Raton, FL.

Hopper, S.D. & Gioia, P. 2004. The Southwest Australian Floristic Region: Evolution and conservation of a global hot spot of biodiversity. Annual Review of Ecology, Evolution, and Systematics, 35, 623–650.

Isbell, R.F. 2002. The Australian Soil Classification, Revised Edition. CSIRO Publishing, Collingwood, Victoria, Australia.

Jangid, K., Whitman, W.B., Condron, L.M., Turner, B.L. & Williams, M.A. 2013. Soil bacterial community succession during long-term ecosystem development. Molecular Ecology, 22, 3415–3424.

Kendrick, G.W., Wyrwoll, K.-H. & Szabo, B.J. 1991. Pliocene-Pleistocene coastal events and history along the western margin of Australia. Quaternary Science Reviews, 10, 419–439.

Kitayama, K. 2005. Comment on “Ecosystem properties and forest decline in contrasting long-term chronosequences”. Science, 308, 633.

Krüger, M., Teste, F.P., Laliberté, E., Lambers, H., Coghlan, M., Zemunik, G. & Bunce, M. 2015. The rise and fall of arbuscular mycorrhizal fungal diversity during ecosystem retrogression. Molecular Ecology, 24, 4912–4930.

Laliberté, E., Grace, J.B., Huston, M.A., Lambers, H., Teste, F.P., Turner, B.L. & Wardle, D.A. 2013. How does pedogenesis drive plant diversity?. Trends in Ecology & Evolution, 28, 331–340.

Laliberté, E., Turner, B.L., Costes, T., Pearse, S.J., Wyrwoll, K.-H., Zemunik, G. & Lambers, H. 2012. Experimental assessment of nutrient limitation along a 2-million-year dune chronosequence in the south-western Australia biodiversity hotspot. Journal of Ecology, 100, 631–642.

Laliberté, E., Zemunik, G. & Turner, B.L. 2014. Environmental filtering explains variation in plant diversity along resource gradients. Science, 345, 1602–1605.

Lambers, H., Clode, P.L., Hawkins, H.-J., Laliberté, E., Oliveira, R.S., Reddell, P., Shane, M.W., Stitt, M. & Weston, P. 2015. Metabolic Adaptations of the Non-Mycotrophic Proteaceae to Soils With Low Phosphorus Availability. In. Annual Plant Reviews Volume 48, pp. 289–335. John Wiley & Sons, Inc.

Loeppert, R.H. & Suarez, D.L. 1996. Carbonate and Gypsum. In. Methods of Soil Analysis, Part 3- Chemical Methods (ed. Sparks, D.L.E.A.), pp. 437–474. Soil Science Society of America Madison, Wisconsin.

McArthur, W.M. 2004. Reference Soils of South-western Australia. Department of Agriculture (WA), Perth, Australia.

McArthur, W.M. & Bettenay, E. 1974. Development and Distribution of Soils of the Swan Coastal Plain, Western Australia. Canberra, Australia.

Peltzer, D.A., Wardle, D.A., Allison, V.J., Baisden, W.T., Bardgett, R.D., Chadwick, O.A., Condron, L.M., Parfitt, R.L., Porder, S., Richardson, S.J., Turner, B.L., Vitousek, P.M., Walker, J. & Walker, L.R. 2010. Understanding ecosystem retrogression. Ecological Monographs, 80, 509–529.

Pinheiro, J.C. & Bates, D.M. 2000. Mixed-Effects Models in S and S-PLUS. Springer, New York, USA.

Playford, P.E., Cockbain, A.E. & Lowe, G.H. 1976. Geology of the Perth Basin, Western Australia; Bulletin 124 of the Geological Survey of Western Australia. Geological Survey of Western Australia, Perth, Australia.

Porder, S. & Chadwick, O.A. 2009. Climate and soil-age constraints on nutrient uplift and retention by plants. Ecology, 90, 623–636.

Purdie, B., Tille, P. & Schoknecht, N. 2004. Soil-landscape mapping in south-Western Australia⍰: an overview of methodology and outputs. Department of Agriculture and Food, Western Australia, Perth, Australia.

Selmants, P.C. & Hart, S.C. 2010. Phosphorus and soil developement: Does the Walker and Syers model apply to semiarid ecosystems?. Ecology, 91, 474–484.

Soil Survey Staff. 1999. Soil Taxonomy: A Basic System of Soil Classification for Making and Interpreting Soil Surveys. United States Department of Agriculture–Natural Resources Conservation Service, Lincoln, NE.

Teste, F.P., Kardol, P., Turner, B.L., Wardle, D.A., Zemunik, G., Renton, M. & Laliberté, E. 2017. Plant-soil feedback and the maintenance of diversity in Mediterranean-climate shrublands. Science, 355, 173–176.

Turner, B.L. & Laliberté, E. 2015. Soil development and nutrient availability along a 2 million-year coastal dune chronosequence under species-rich Mediterranean shrubland in southwestern Australia. Ecosystems, 18, 287–309.

Turner, B.L. & Romero, T.E. 2009. Short-term changes in extractable inorganic nutrients during storage of tropical rain forest soils. Soil Science Society of America Journal, 73, 1972–1979.

Vitousek, P. & Chadwick, O. 2013a. Pedogenic thresholds and soil process domains in basalt-derived soils. Ecosystems, 16, 1379–1395.

Vitousek, P.M. 2004. Nutrient Cycling and Limitation. Princeton University Press, Princeton, New Jersey.

Vitousek, P.M. & Chadwick, O.A. 2013b. Pedogenic thresholds and soil process domains in basalt-derived soils. Ecosystems, 16, 1379–1395.

Vitousek, P.M. & Farrington, H. 1997. Nutrient limitation and soil development: Experimental test of a biogeochemical theory. Biogeochemistry, 37, 63–75.

Walker, T.W. & Adams, A.F.R. 1958. Studies on soil organic matter: I. Influence of phosphorus content of parent materials on accumulations of carbon, nitrogen, sulfur, and organic phosphorus in grassland soils. Soil Science, 85, 307–318.

Walker, T.W. & Syers, J.K. 1976. The fate of phosphorus during pedogenesis. Geoderma, 15, 1–19.

Wardle, D.A., Walker, L.R. & Bardgett, R.D. 2004. Ecosystem properties and forest decline in contrasting long-term chronosequences. Science, 305, 509–513.

Williamson, W.M., Wardle, D.A. & Yeates, G.W. 2005. Changes in soil microbial and nematode communities during ecosystem decline across a long-term chronosequence. Soil Biology and Biochemistry, 37, 1289–1301.

Wyrwoll, K.-H., Turner, B.L. & Findlater, P. 2014. On the origins, geomorphology and soils or the sandplains of south-western Australia. In. Plant Life on the Sandplains in Southwest Australia, a Global Biodiversity Hotspot (ed. Lambers, H.). University of Western Australia, Crawley, Australia.

Zemunik, G., Turner, B.L., Lambers, H. & Laliberté, E. 2015. Diversity of plant nutrient-acquisition strategies increases during long-term ecosystem development. Nature Plants, 1.

Zemunik, G., Turner, B.L., Lambers, H. & Laliberté, E. 2016. Increasing plant species diversity and extreme species turnover accompany declining soil fertility along a long-term chronosequence in a biodiversity hotspot. Journal of Ecology, 104, 792–805.

